# ATF4 alleviates doxorubicin-induced cardiomyopathy through H_2_S-mediated antioxidation

**DOI:** 10.1101/2025.09.03.674119

**Authors:** Shuting Xu, Yan Shi, Xiaoshuai Zhao, Xuhui Chen, Ying Liu, Fan Zhang, Fangchi Yu, Linhao Ruan, Chaolong Wang, Xuejun Jiang, Xiaoding Wang, Guangyu Zhang

## Abstract

**Background:** Doxorubicin (DOX) remains a cornerstone in the treatment of various cancers. However, its clinical utilization is significantly hampered by dose-dependent cardiotoxicity. The generation of reactive oxygen species (ROS) constitutes the central component of the pathogenesis of DOX-induced cardiotoxicity. Activating transcription factor 4 (ATF4) has been demonstrated to exert a cardioprotective effect and augment cardiac antioxidative capacity in settings of heart failure. However, the role of ATF4 in DOX-induced cardiomyopathy (DIC) remains unknown.

**Methods:** To explore the role of ATF4 in DOX-induced cardiomyopathy, cardiac-specific ATF4 conditional heterozygous mice and AAV9 mediated ATF4 overexpression mouse models were utilized. Cardiac function was assessed by echocardiography. The upstream regulator and downstream mediator of ATF4 were evaluated using RNA-seq analysis and further verified using ChIP assay and luciferase reporter assay.

**Results:** We found a substantial decrease in ATF4 expression levels in the heart of DIC mice. ATF4^+/-^ mice exhibited a higher degree of susceptibility to DOX-induced cardiotoxicity in comparison with ATF4^flox/flox^ mice, as evidenced by the manifestation of more severe cardiac dysfunction and a significantly earlier mortality rate. In contrast, cardiacc-specific overexpression ATF4 by AAV9 confers robust cardioprotection against DOX-induced cardiomyopathy. Mechanistically, we identified the upstream regulator of ATF4 as KLF16, which was significantly suppressed during DOX treatment. Further, the decrease of ATF4 led to a reduction in cystathionine γ-lyase (CSE) transcription and hydrogen sulfide (H_2_S) production in the context of DOX-induced cardiotoxicity. ChIP and luciferase reporter assays revealed that ATF4 functioned as the transcription factor of the CSE gene, which is a key enzyme in the synthesis of H_2_S to counteract oxidative stress. Consistently, ROS scavengers or H_2_S donors was shown to mitigate the consequences of ATF4 deficiency. In contrast, the ectopic expression of ATF4 mitigated oxidative stress and apoptosis in DOX-induced cardiotoxicity, both *in vivo* and *in vitro*.

**Conclusions:** Our study revealed a novel function of ATF4 in counteracting oxidative stress in DOX cardiotoxicity by promoting the transcription of CSE. ATF4 may represent a promising therapeutic target for the treatment of DOX-induced cardiomyopathy.

## 1. Introduction

Doxorubicin (DOX), a potent anthracycline chemotherapeutic agent, remains a cornerstone in the treatment of various malignancies, including breast cancer, lymphomas, and sarcomas [1]. However, its clinical utilization is significantly hampered by dose-dependent cardiotoxicity, leading to DOX-induced cardiomyopathy (DIC). This condition manifests as irreversible myocardial damage, left ventricular dysfunction, and congestive heart failure, with mortality rates exceeding 50% within two years of diagnosis [2]. The observed cumulative toxicity of DOX, particularly at doses exceeding 450 mg/m², underscores the urgent need to understand its pathogenic mechanisms and develop cardioprotective strategies [3]. The generation of reactive oxygen species (ROS) is a central component of DOX cardiotoxicity, as it induces mitochondrial dysfunction, disrupts calcium homeostasis, and activates pro-cell death pathways [4]. Recent studies have also emphasized the significance of ferroptosis, an iron-dependent form of regulated cell death, in DIC [5, 6]. Additionally, DOX has been shown to intercalate into nuclear DNA, resulting in the formation of double-strand breaks and the impairment of topoisomerase IIβ activity. This, in turn, has been shown to trigger cardiomyocyte senescence [7, 8]. Despite these processes in our understanding of DOX cardiotoxicity, current preventive approaches, including liposomal DOX formulations and dexrazoxane treatment, offer limited cardioprotection [9, 10].

ATF4 (Activating Transcription Factor 4), a master transcription factor governing cell survival and metabolism, occupies a central position in orchestrating adaptive and maladaptive responses to diverse stressors. Mechanistically, the activation of ATF4 is subject to stringent regulation by the integrated stress response (ISR), wherein four upstream kinases (PERK, GCN2, PKR, and HRI) phosphorylate the α-subunit of eukaryotic initiation factor 2 (eIF2α) [11]. This phosphorylation event has been shown to attenuate global protein translation while selectively enhancing ATF4 mRNA translation, thereby enabling rapid transcriptional reprogramming under stress conditions. ATF4 serves as a central regulatory hub that orchestrates cellular responses to various stressors, including endoplasmic reticulum (ER) stress, nutrient deprivation, viral infection, double-stranded RNA accumulation, and heme deficiency [12]. Upon activation, ATF4 translocates to the nucleus and binds to AAREs or CAREs in target gene promoters. This transcriptional control regulates a network of genes involved in redox homeostasis, amino acid metabolism, and unfolded stress response [13].

Our previous studies demonstrated that ATF4 plays a cardioprotective role and enhances cardiac antioxidative capacity under the condition of pressure overload and pathological cardiac remodeling [14]. However, the function of ATF4 in DOX cardiotoxicity remains unknown. It has been demonstrated that DOX triggers ER stress, oxidative damage, and iron dysregulation, all of which are known activators of the ISR-ATF4 axis. Here we set out to investigate the role of ATF4 in DOX cardiotoxicity and explore the underlying molecular mechanisms.

## 2. Materials and methods

### Animals

All mouse procedures were approved by the Institutional Animal Care and Ethics Committee of the Tongji Medical School, Huazhong University of Science and Technology (HUST, Wuhan, China). Mice were on the C57BL/6J background and maintained in a 12:12 hours light:dark cycle in temperature-controlled rooms with unlimited access to water and chow food. To achieve cardiac-specific deletion of ATF4, we generated ATF4 heterozygous mice (Atf4 flox/+; Myh6-Cre). Genotyping primers are listed in Supplemental Table S1.

### Doxorubicin treatment *in vivo*

To induce acute DOX cardiotoxicity, mice were injected with a single dose of DOX (15 mg/kg) or an equal volume of normal saline (NS) via intraperitoneal injection(IP). Mice were weighed daily and euthanized on day 7 post-DOX injection.

### Echocardiography

Cardiac function was evaluated in conscious, unconstrained mice using echocardiography (Visual Sonics, #Vevo 2100, MS400C probe) as previously described [14]. M-mode images were captured in the short-axis view at the level of the papillary muscles and analyzed to calculate various parameters. Heart rate was recorded. Left ventricular internal diameters (in mm) at end diastole (LVID, diastolic) and end systole (LVID, systolic) were determined using M-mode recordings. Fractional shortening and ejection fraction were calculated as a percentage.

### Chemicals

Doxorubicin (DOX, APEXBIO, #A1832) was dissolved in H_2_O to make 50mM stock solution, which was used *in vitro* at concentration of 1 μM or 2 μM. DPI (MCE, #HY-100965) was dissolved in DMSO to make 10 mM stock and used at 500 nM. Sodium hydrosulfide (NaHS, Adamas, #16721-80-5) was dissolved in water to make 5M stock and final concentration was 1.6mM.

### Cell culture and siRNA treatment

H9c2 cells, derived from embryonic rat myocardium, were purchased from the Procell (#CL-0089). Cells were cultured in high-glucose DMEM medium, supplemented with 10% fetal bovine serum and 1 % antibiotic and antimycotic solution. Similarly, HEK293 cells were cultured under the same conditions. AC16 cells were purchased from the Procell (#CL-0790), cultured in DMEM/F12 medium, supplemented with 10% fetal bovine serum, 1 % antibiotic, and antimycotic solution. Hela cells were purchased from the Procell (#CL-0101), cultured in MEM medium, supplemented with 10% fetal bovine serum, 1 % antibiotic, and antimycotic solution. All cell lines were maintained in a humidified incubator at 37°C with 5% CO₂ and were passaged every 2–3 days. Routine mycoplasma contamination testing was performed to ensure cell health. Small interfering RNAs (siRNAs) targeting ATF4 were purchased from Sigma (#SASI_Rn01_00103280, #SASI_Rn01_00103284). siRNAs were dissolved in Opti-MEM reduced-serum medium (Thermo Fisher Scientific, #31985070) at a concentration of 37 µM. For transfection, approximately 80 pmol of siRNA was used per well in 6-well plates, employing the BOLG-RNA™ reagent (#BOLG101, Biology). Following a 6-hour incubation period, cells were replenished with fresh culture medium to support continued growth and expression analysis.

### Knockout cell lines generation

ATF4^KO^ H9c2 cells were generated using the CRISPR/Cas9 system. Briefly, three single-guide RNAs (sgRNAs: sgRNA1: TCTCTTAGATGACTATCTGG, sgRNA2: TTGTCGCTGGAGAACCCATG, sgRNA3: TCATGGGTTCTCCAGCGACA) were designed targeting exon 3 of the *ATF4* gene. For exon knockout, H9c2 were electroporated with Cas9 protein and sgRNA. Single clones were isolated and enriched. Colonies were genotyped by PCR and sequencing to validate successful ATF4 knockout. Two single-guide RNAs (sgRNAs: sgRNA1: GGGTCGCCCGCGATGCACCA, sgRNA2: CGGGAGCCGTGGTGCATCGC) were designed targeting exon 1 of the *KLF16* gene.

### Neonatal rat ventricular myocyte (NRVM) isolation and treatment

Neonatal rat ventricular myocytes (NRVMs) were isolated from the ventricles of 1-day-old Sprague-Dawley rats using a cardiomyocyte isolation kit (Cellutron, #NC-6031), following previously established protocols [15]. To eliminate neonatal fibroblasts, NRVMs were pre-plated for 2 hours. Subsequently, the cells were plated at a density of 1,250 cells/mm² in plating medium, which consisted of a 3:1 ratio of DMEM to M199, supplemented with 5% fetal bovine serum (FBS), 10% horse serum, 1% penicillin/streptomycin, and 100 µM bromodeoxyuridine (BrdU). After 24 hours, the NRVMs were transferred to a serum-reduced medium (DMEM/M199, 3:1 ratio, with 1% FBS, 1% penicillin/streptomycin, and 100 µM BrdU). Following an additional 24 hours, the cells were switched to serum-free medium (DMEM/M199, 3:1 ratio). NRVMs were then subjected to various experimental treatments, such as siRNA-mediated knockdown and adenovirus infection.

### RNA isolation, reverse transcription and quantitative PCR

Total RNA from cardiac tissues and cells was extracted with an Aurum Total RNA Fatty and Fibrous Tissue kit (Bio-Rad, 732-6870) and a Quick-RNA Microprep kit (Zymo Research, R1051), respectively. A total of 250 ng RNA was used for reverse transcription (Takara, RR036A), and cDNA was diluted 10-fold by ddH_2_O. For each sample, 2 µL of cDNA was used for quantitative PCR to determine relative mRNA levels relative to 18s rRNA using a Light Cycler machine (Roche) and the SYBR green reagent (Bio-Rad, 1725125). All primer sequences are listed in Table I of the online Data Supplement.

### Cell death assay

Cell viability of H9c2 cells were assessed using the CCK8 assay. Briefly, CCK8 was diluted in DMEM medium with 10% FBS at 1:10 as working solution. After treated with DOX, the culture medium of H9c2 cells was replaced with CCK8 working solution. Following a 30-minute incubation at 37°C, the absorbance was measured at 450 nm and cell viability was calculated as indicated. NRVMs cell death was assayed by measuring LDH release using a CytoTox96 non-radioactive cytotoxicity assay kit (Promega, #G1780) according to the manufacturer’s instructions.

### SYTOX Green /Hoechst double staining

SYTOX Green (Thermo Fisher Scientific, #S7020) and Hoechst (Thermo Fisher Scientific, #H3570) were diluted in DMEM medium with 10% FBS at 1:1000 as working solution. After treated with DOX, the culture medium of H9c2 cells was changed to SYTOX Green work solution. After incubating at 37°C for 30 minutes, cells were washed for 3 times before imaging with a fluorescence microscope (Nexcope, NIB610-FL).

### Detection of intracellular H_2_S

SF-7AM probe (APEXBIO, # C3504) was used to detect intracellular H_2_S level. Briefly, H9c2 cells were exposed to 2.5 μM SF-7AM for 30 minutes after treated with or without DOX. After washing, fluorescent images of cells were captured with a fluorescence microscope (Nexcope, NIB610-FL), and cellular intensity of fluorescence was quantified using ImageJ software.

### Measurement of intracellular ROS

The levels of intracellular ROS were determined by spectrophotometry using DCFH-DA (Beyotime, #S0033S). Cells were washed with PBS and incubated with 10 μM fluorescence dyes (final concentration) for 30 minutes at 37°C in the dark. Next, cells were washed three times with PBS, and the fluorescence intensity was determined.

### Immunoblotting

Total proteins were extracted from cultured cardiomyocytes or cardiac tissues using RIPA lysis buffer supplemented with protease and phosphatase inhibitors (Thermo Fisher Scientific, #88669). Protein concentration was measured using a BCA kit (Servicebio, #G2026-200T) according to the manufacturer’s instructions. For SDS-PAGE analysis, concentration-normalized samples were prepared by mixing the lysates with 2× loading buffer and boiling at 99℃ for 15 minutes. Samples were then loaded onto 10% or 12.5% SDS-PAGE gels and transferred onto nitrocellulose membranes (Bio-Rad, #1704157). The membranes were blocked for 1 hour at room temperature with 5% BSA and subsequently incubated with primary antibodies overnight at 4°C. After washing, the membranes were incubated with fluorescent dye-labeled secondary antibodies at room temperature for 1 hour. Following extensive washing with TBS-Tween 20, the blots were scanned using a Tanon 5200 imager. The primary antibodies used were as follows: GAPDH (Fitzgerald, #10R-G109A), ATF4 (Cell Signaling Technology, #11815), p-eIF2α (Thermo Fisher Scientific, #44-728G), Gamma Cystathionase (CSE, Proteintech, #12217-1-AP), Cleaved-Caspase 3 (Cell Signaling Technology, #9664S), Caspase 3 (Cell Signaling Technology, #9662), KLF16 (Immunoway, #YN0327), GPX4 (Abcam, #ab125066), GSDMD (Abcam, #ab219800), LC3B (ablconal, #A19665), MLKL (Sigma, #MABC604), Phospho-MLKL (Ser345) (Cell Signaling Technology, # 37333), 4-Hydroxynonenal Antibody (4-HNE, R&D system, # MAB3249). The secondary antibodies used were HRP-conjugated AffiniPure goat anti-rabbit IgG (H+L) (BOSTER, #BA1055) and HRP-conjugated AffiniPure goat anti-mouse IgG (H+L) (BOSTER, #BA1051).

### Adenovirus production and infection

ATF4 fusion gene was used for adenovirus generation with Adeno-X adenoviral system 3 (Takara, #632269). Adenoviruses expressing blank vector (NC) were used as controls.

### Adeno-associated virus serotype 9 production

Mouse ATF4 cDNA was amplified from a mouse heart cDNA library and cloned into the pAV-cTNT-P2A-GFP construct. This plasmid contains a chicken troponin T promoter to enhance gene expression in a cardiac-specific manner. AAV9 virus packaging was achieved in HEK293Tcells, purified and concentrated by gradient centrifugation (WZ Biosciences Inc). Mice were treated with AAV9 4E10^10 particles/mouse before DOX injection with AAV9-GFP as a negative control.

### Histology and immunostaining

Mouse hearts were harvested and fixed in 10% neutralized formalin for 48 hours at 4°C. Paraffin sections (5-μm thickness) were prepared. Hematoxylin and eosin (H&E) and Masson’s trichrome staining were used to assess heart morphology and myocardial fibrosis. Histological images with the least processing artifact and denoting relatively similar cell numbers for each antibody or similar anatomy were chosen as representative images.

### Chromatin immunoprecipitation assay (ChIP)

ChIP was performed using a ChIP assay kit (Beyotime, #P2078). Briefly, H9c2 cells were first infected by adenoviruses expressing ATF4. Cells were cross-linked by 37% formaldehyde, lysed with the SDS Lysis Buffer, and subjected to sonication to shear DNA. Cross-linked protein/DNA was precipitated using anti-flag tag antibody (Beyotime, #AF2852) with normal mouse IgG antibody as a control. PCR was then conducted to compare ATF4-CSE promoter binding.

### Luciferase reporter assay

The promotor region of CSE, including the transcription factor ATF4 binding site (AAGATGTAAA), was amplified from the genomic DNA of rat cells. Reportor construct was generated by linking the CSE promotor into the PGL3-basic vector. Hela cells was infected with ATF4 overexpression adenovirus and next transfected with the indicated luciferase constructs and internal control plasmid phRL-TK. Firefly luciferase and Renilla luciferase activity was measured according to the manual of the Dual-luciferase reporter assay kit (Beyotime cat # RG027).

### Statistical analyses

Data are represented as the mean±SEM. Two-tailed Student’s t-test was employed to compare the differences between two groups. For multiple group comparisons with one variable, one-way ANOVA was conducted, followed by Tukey multiple comparisons test. For multiple group comparisons with two or more variables, two-way ANOVA was conducted, followed by either Tukey or Sidak multiple comparisons test. A P value of <0.05 was considered statistically significant. Statistical analyses were performed using GraphPad Prism software 9.3.0

## 3. Results

### 3.1. ATF4 is downregulated in DOX-induced cardiomyopathy

We searched the GEO database and found a dataset in which the cardiac spheroid samples were treated with DMSO, unformulated DOX at 5 μM, or SPEDOX-6 at 5 μM DOX equivalent dose for 48 hours (GSE235470). We found that the mRNA levels of ATF4 were significantly downregulated, and the transcription factor binding pathway is enriched substantially (Fig. 1A, S1A). We next decided to evaluate the expression of ATF4 in cardiomyocytes under DOX treatment. To explore the expression of ATF4 *in vivo*, a mouse model of DOX-induced cardiomyopathy was generated by i.p. injection of a dose of 15 mg/kg DOX (Fig. S1B). Morphological examination revealed that the heart in the DOX treatment group was smaller than that in the control group (Fig. 1B), and the heart weight to tibia length ratio (HW/TL) significantly reduced in the DOX treatment group despite a minor decrease of the heart weight to body weight ratio (HW/BW) (Fig. S1C). Echocardiography demonstrated a downward trend in cardiac systolic function in the DOX treatment group, as evidenced by significantly decreased left ventricular ejection fraction (EF) and fractional shortening (FS) (Fig. 1C, S1D and E). H&E staining and Masson’s trichrome staining showed that DOX treatment exacerbated cardiac fibrosis relative to the saline treatment (Fig. S1F). These results indicate that we successfully established a DOX-induced cardiomyopathy model. Notably, the expression of ATF4 was substantially downregulated in DOX-treated hearts as shown by RT-PCR (Fig. 1D). *In vitro*, a significant decrease of ATF4 expression was observed in DOX-treated H9c2 cardiomyocytes, as demonstrated by RT-PCR and immunoblot (Fig. 1E-G). Similarly, the expression of ATF4 was reduced in DOX-treated AC16 cells and NRVMs, respectively (Fig. 1H-J, S1G and H).

**Figure 1.**
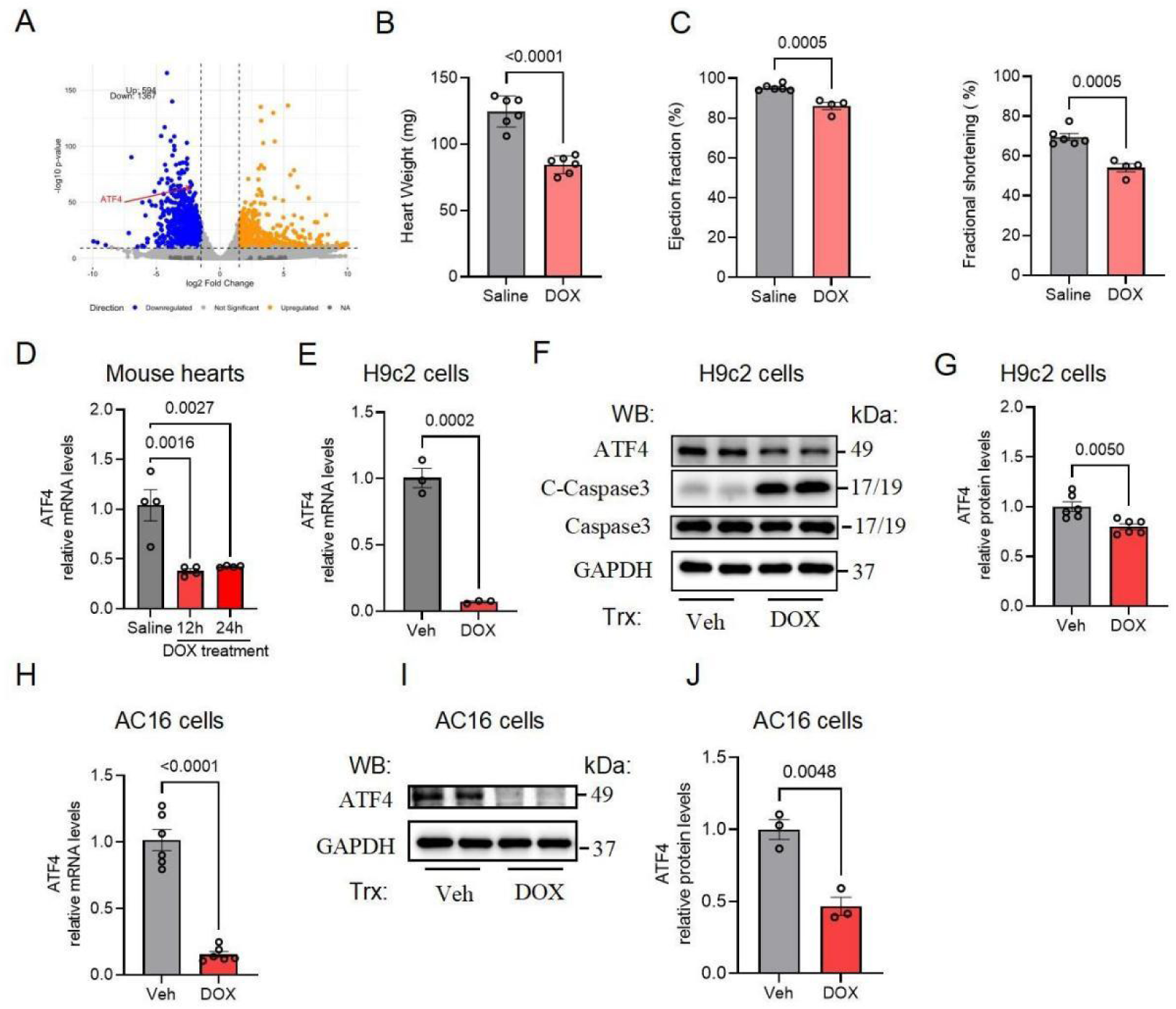
ATF4 is downregulated in DOX-induced cardiomyopathy. **(A)** Volcano plot detailing significant gene expression changes for DOX vs. Saline, with upregulated genes in orange and downregulated genes in blue. **(B)** DOX treatment reduced heart weight (n = 6 mice per group). **(C)** DOX led to cardiac dysfunction 1 week after treatment, as revealed by decreases in left ventricular ejection fraction (EF) and left ventricular fractional shortening (FS) (n = 6 for Saline; n = 4 for DOX). **(D)** RT-qPCR revealed that DOX treatment led to a decrease in ATF4 mRNA levels in the mouse heart (n = 4 mice per group). **(E)** RT-qPCR revealed that DOX treatment led to a decrease in ATF4 mRNA levels in the H9c2 cells (n = 3 per group). **(F)** Western blot revealed that DOX treatment led to a decrease in the ATF4 protein levels and a increase in the cleaved-caspase 3 protein level in H9c2 cells (n = 6 per group). **(G)** Quantification of (F). **(H)** RT-qPCR revealed that DOX treatment led to a decrease in ATF4 mRNA levels in the AC16 cells (n = 6 per group). **(I)** Western blot revealed that DOX treatment led to the decrease of ATF4 protein level in the AC16 cells (n = 3 per group). **(J)** Quantification of (I). Data are represented as mean ± SEM.

### 3.2. ATF4 knockdown exacerbates DOX-induced cardiomyopathy *in vivo*

To determine the role of ATF4 *in vivo*, we crossed conditional ATF4^F/F^ animals with cardiomyocyte-specific Cre transgenic mice (ɑMHC-Cre). According to our previous publication, homozygous knockout mice (ATF4 cKO) manifest appreciable differences at the level of ROS production in the heart at baseline, as validated by 4HNE staining of the heart tissue [14]. We then generated cardiomyocyte-specific heterozygous mice (ATF4 Het) (Fig. 2A). We observed that ATF4 mRNA level was significantly decreased in the hearts of heterozygous mice compared to the control mice (Fig. 2B). We firstly checked the ROS production levels in ATF4 Het mice utilizing 4HNE staining and did not find significant change compared with the control mice (Fig. S2A and B). Cardiac function analysis of ATF4 Het mice did not reveal significant changes compared to the control mice (Fig. S2C and D). We then established the DIC model in ATF4 Het mice. We firstly estimated the survival rates of these mice and found that ATF4 heterozygous deletion significantly accelerated the death of DOX-treated mice (Fig. 2C). We also found that DOX treatment led to a conceivable decline of EF and FS, as assayed by echocardiography assessment (Fig. 2D, E, S2E). Morphological examination revealed that the size of heart in the DOX treatment group was smaller than that in the control group (Fig. S2F). In addition, the fibrotic state of hearts was investigated by HE and Masson’s trichrome staining. Our analysis revealed that DOX-induced cardiac fibrosis was aggravated by ATF4 heterozygous deletion (Fig. 2F and G). Taken together, these *in vivo* data indicate that ATF4 reduction promotes DOX-induced cardiac injury and exacerbates the progression of DIC.

**Figure 2.**
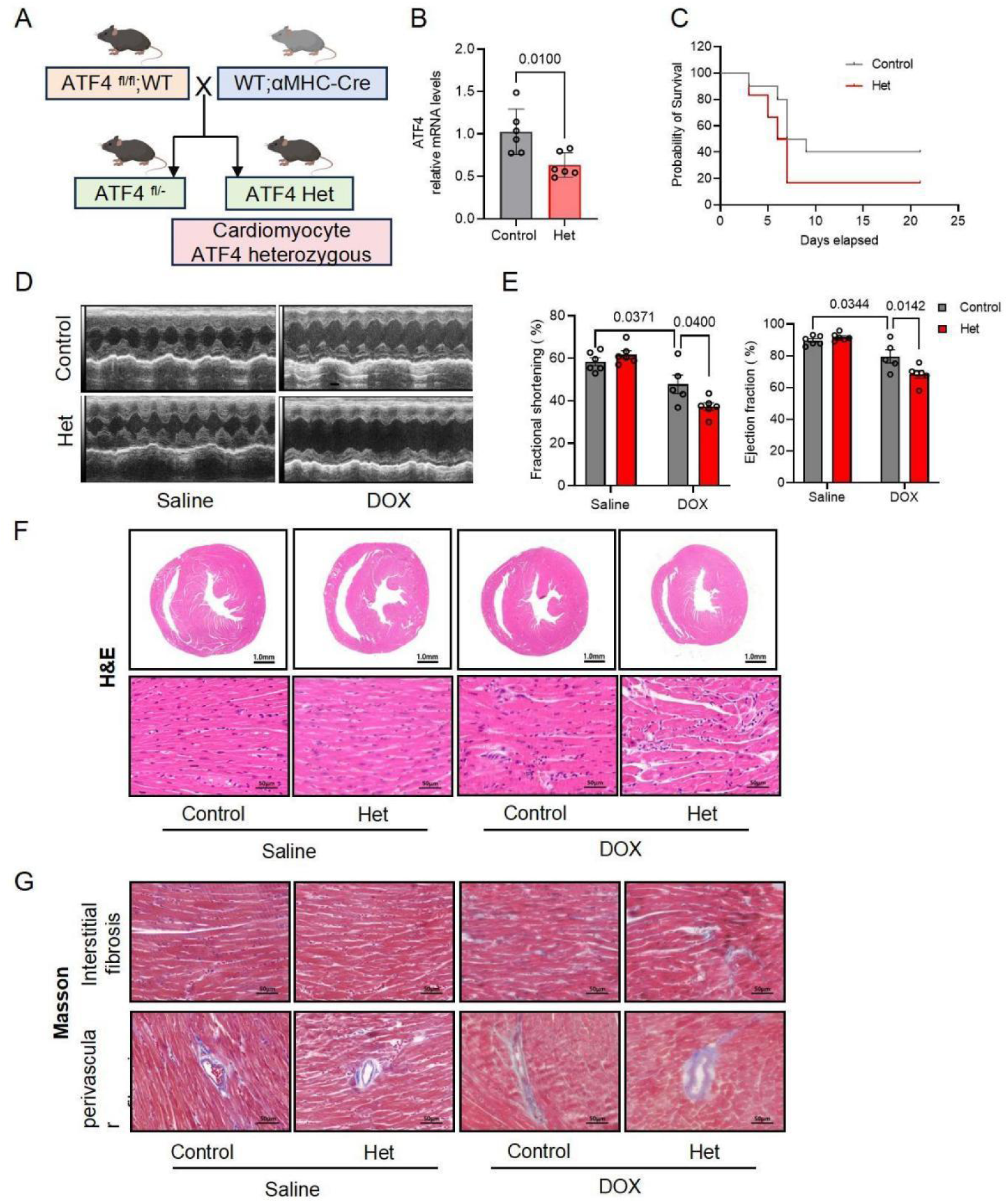
ATF4 knockdown exacerbates DOX-induced cardiomyopathy in vivo. **(A)** Schematic of the cardiac-specific ATF4 conditional heterozygous mouse model (Het). **(B)** Validation of ATF4 conditional heterozygous mouse. RT-qPCR was conducted, and the ATF4 relative mRNA level was quantified (n = 6). **(C)** Kaplan-Meier survival curves of the indicated mice treated with DOX. (n = 10 per group). **(D)** Control ATF4^fl/-^ (Control) and ATF4 cardiomyocyte-specific conditional knockdown (Het) mice were used for echocardiography measurements 1 week after DOX treatment. Representative M-mode images are shown. **(E)** ATF4 knockdown led to deterioration of cardiac function 1 week after DOX treatment, as revealed by decreases in EF and FS (n = 6 for Saline, Control; n = 6 for Saline, Het; n = 5 for DOX, Control; n = 6 for DOX, Het). **(F)** Representative Hematoxylin & Eosin (H&E) staining of heart (scale bar = 1mm/50μm) in four groups of mice. **(G)** Representative Masson’s Trichrome staining of heart (scale bar = 50 μ m) in four groups of mice. Data are represented as mean ± SEM.

### 3.3. ATF4 overexpression alleviates DOX-induced cardiomyopathy *in vivo*

We next asked whether ATF4 overexpression may protect the heart from DIC. Wild-type C57BL/6J male mice were infected with adeno-associated virus 9 (AAV9) carrying ATF4 (AAV9-ATF4) to overexpress ATF4 or AAV9-GFP as negative control by jugular vein injection. Western blot analysis confirmed that ATF4 expression was upregulated in AAV9-ATF4 group versus the AAV9-GFP group (Fig. 3A). Then, these mice were injected with DOX or normal saline (Fig. 3B). We found that ATF4 overexpression significantly improved EF and FS compared with those in the control group after treated with DOX (Fig. 3C, D, S2G). In addition, H&E and Masson’s trichrome staining demonstrated that ATF4 overexpression also alleviated DOX-induced cardiac fibrosis relative to the control treatment (Fig. 3E and F). Taken together, these *in vivo* findings suggest that ATF4 overexpression alleviates DOX-induced cardiac injury and mitigates the progression of DIC.

**Figure 3.**
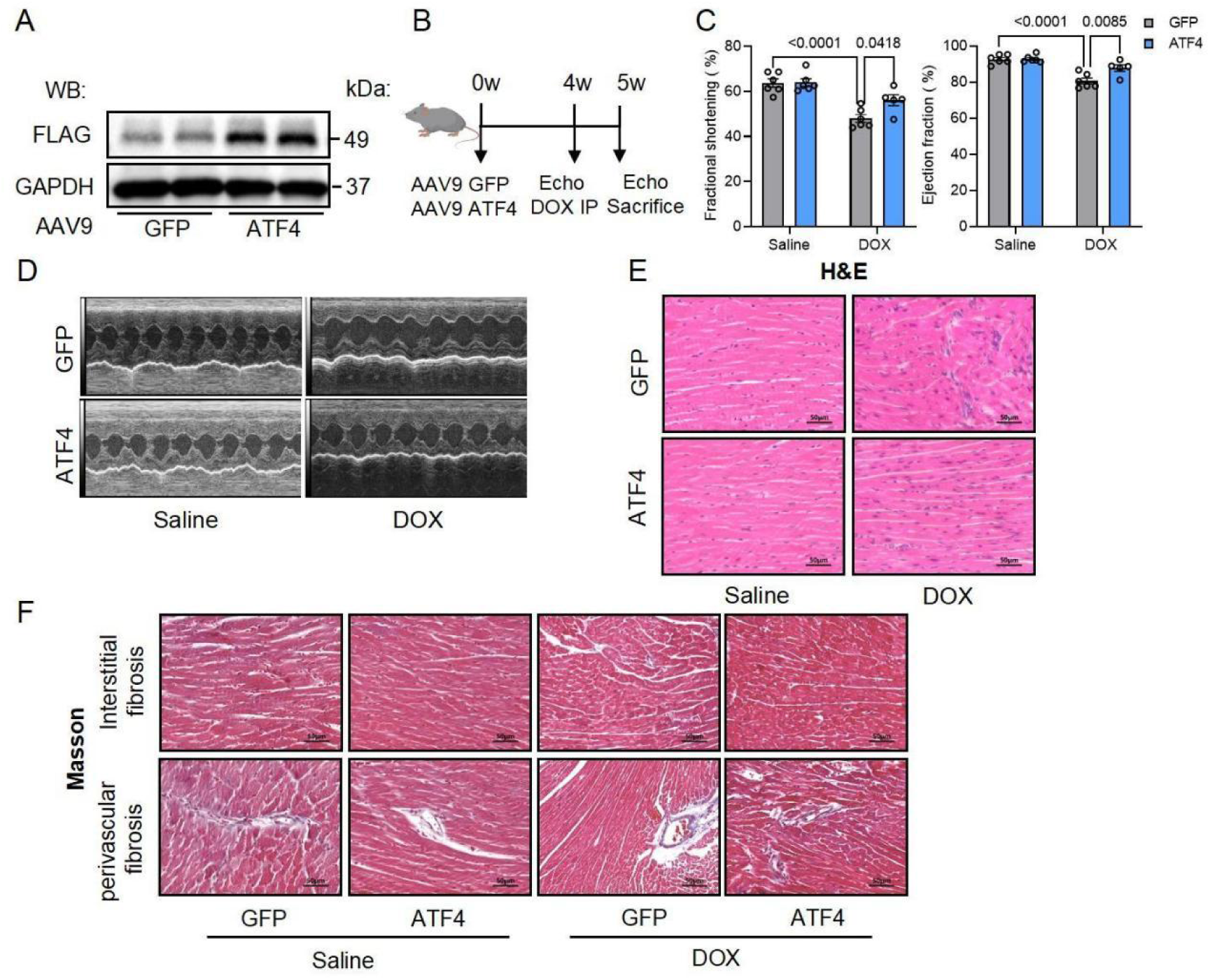
ATF4 overexpression alleviates DOX-induced cardiomyopathy in vivo. **(A)** Validation of ATF4 overexpression. Western blot was conducted and flag relative protein level was shown. **(B)** Scheme of AAV9 injection and DOX administration. **(C)** ATF4 overexpression led to improvement of cardiac function 1 week after after DOX treatment, as revealed by decreases in EF and FS (n = 6 for Saline, GFP; n = 6 for Saline, ATF4; n = 6 for DOX, GFP; n = 5 for DOX, ATF4).**(D)** Mice injected with GFP-AAV9 and ATF4 AAV9 were used for echocardiography measurements 1 week after DOX treatment. Representative M-mode images are shown. **(E)** Representative Hematoxylin & Eosin(H&E) staining of heart (scale bar = 1.0 mm/50μm) in four groups of mice. **(F)** Representative Masson’s Trichrome staining of heart (scale bar = 50μm) in four groups of mice. Data are represented as mean ± SEM.

### 3.4. ATF4 ameliorates DOX-mediated cardiomyocyte damage *in vitro*

To evaluate the effects of ATF4 deficiency in DOX-induced cardiomyocyte damage, H9c2 cells, AC16 cells, and NRVMs were employed. Using CRISPR-Cas9-mediated genome engineering system, we generated an ATF4 knockout (ATF4^KO^) H9c2 cell line (Fig. S3A). Western blot analysis confirmed that ATF4 protein level significantly decreased in ATF4^KO^ cells compared to the control cells (Fig. 4A). A marked decrease in cell survival ratio was observed in ATF4^KO^ cells utilizing CCK-8 assay after treated with DOX compared with ATF^WT^ cells (Fig. 4B). Similarly, our phase-contrast microscopic analysis demonstrated that ATF4 deficiency exacerbated DOX-mediated cardiomyocyte death in H9c2 cell as indicated by the Sytox Green staining (Fig. 4C and D). To confirm this phenotype, ATF4 expression was reduced in AC16 cells and NRVMs by transfecting siRNA transfection. RT-qPCR and western blot confirmed that ATF4 mRNA and protein levels significantly decreased in AC16 cells transfected ATF4 siRNA compared with those transfected control siRNA (Fig. S3B-D). We found that ATF4 knockdown exacerbate DOX-mediated cardiomyocyte damage in AC16 cells and NRVMs as indicated by CCK-8 and LDH release assay (Fig. 4E and F). To further examine the effect of ATF4 in DOX-induced cardiomyocyte injury, we transfected adeno virus carrying ATF4 (adATF4) to overexpress ATF4 in H9c2 cells. Western blot analysis confirmed that ATF4 expression was upregulated in adATF4 infected H9c2 cells versus the control cells (Fig. 4G). CCK-8 assay demonstrated that cell viability increased in the ATF4 overexpressed cells compared to the control cells after treated with DOX (Fig. 4H). Similarly, Sytox Green staining showed that cell death significantly decreased in the ATF4 overexpressed cells compared to the control cells after treated with DOX (Fig. 4I and J). To further confirm this phenotype, both AC16 cells and NRVMs were infected with adATF4. CCK-8 and LDH release assay also showed that cell death decreased in the ATF4 overexpressed AC16 cells and NRVMs compared to the control cells (Fig. 4K and L).

**Figure 4.**
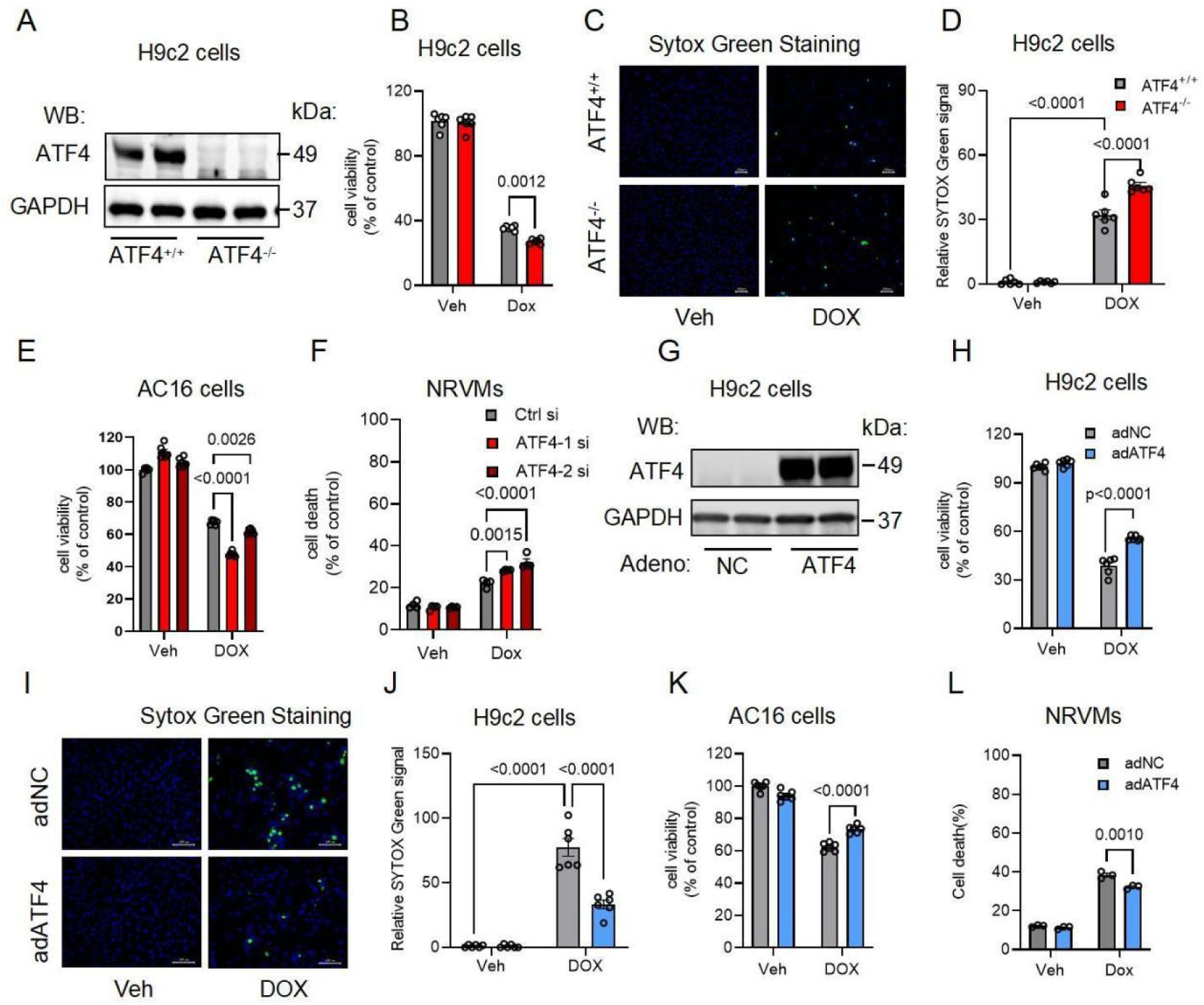
ATF4 regulates DOX-mediated cardiomyocyte damage in vitro. **(A)** Validation of ATF4 knockout H9c2 cells. Western blot was conducted, and the ATF4 relative protein level was shown. **(B)** Cell viability was detected in ATF4^+/+^ and ATF4^−/−^ H9c2 cells respectively, after being treated with DOX utilizing the CCK8 assay (n = 6 per group). **(C)** Representative Sytox green staining images were shown. **(D)** Quantification of (C). **(E)** Cell viability was detected in control and ATF4 knockdown AC16 cells respectively, after being treated with DOX utilizing the CCK8 assay (n = 6 per group). **(F)** Cell death was detected in control and ATF4 knockdown NRVMs respectively, after being treated with DOX utilizing the LDH assay (n = 4 per group). **(G)** Validation of ATF4 overexpression in H9c2 cells. Western blot was conducted, and the ATF4 relative protein level was shown. **(H)** Cell viability was detected in control and ATF4 overexpressed H9c2 cells respectively, after being treated with DOX utilizing the CCK8 assay (n = 6 per group). **(I)** Representative Sytox green staining images were shown. **(J)** Quantification of (I). **(K)** Cell viability was detected in control and ATF4 overexpressed AC16 cells respectively, after being treated with DOX utilizing the CCK8 assay (n = 6 per group). **(L)** Cell death was detected in control and ATF4 overexpressed NRVMs respectively, after being treated with DOX utilizing the LDH assay (n = 3 per group). Data are represented as mean ± SEM.

### 3.5. ATF4 is governed by KLF16 instead of ISR in DIC

As ATF4 is the main response factor of integrated stress response (ISR), we next evaluate the know upstream regulator of ATF4–eIF2α. Unexpectedly, the phosphorylation level of eIF2α was significantly increased instead of decreased (Fig. S4A and B). To identify the upstream regulator of ATF4 in the context of DOX-induced cardiomyocyte damage, we searched all potential transcription factors (TFs) of ATF4 including Krüppel-like factors (KLFs) which belong to the zinc finger-containing TFs. RT-qPCR showed that KLF16 significantly decreased in DIC mice while other KLFs didn’t change (Fig. 5A, S4C). Then we confirmed this in DOX treated H9c2 cells with QPCR and western blotting. We found that KLF16 significantly decreased compared with the control group both at the mRNA and protein level (Fig. 5B-D). We next knocked out KLF16 in H9c2 cells utilizing CRISPR-Cas9-mediated genome engineering system. We found that ATF4 protein level significantly decreased in KLF16 deficiency cells (Fig. 5E and F). After treated with DOX, we found that KLF16 deficiency aggravated DOX-induced cardiomyocyte damage utilizing CCK-8 assay (Fig. 5G). On the other hand, we transferred the KLF16 plasmid to overexpress KLF16 in 293T cells and H9c2 cells. We found that KLF16 overexpression increased the ATF4 protein level (Fig. 5H and I). We then treated KLF16^OE^ H9c2 cells with DOX and found that KLF16 overexpression alleviated DOX-induced cardiomyocyte damage utilizing CCK-8 assay (Fig. 5J, S4D). These results together indicated that KLF16 is the upstream regulator of ATF4 in DOX-induced cardiomyocyte damage instead of the canonical ISR-eIF2α pathway.

**Figure 5.**
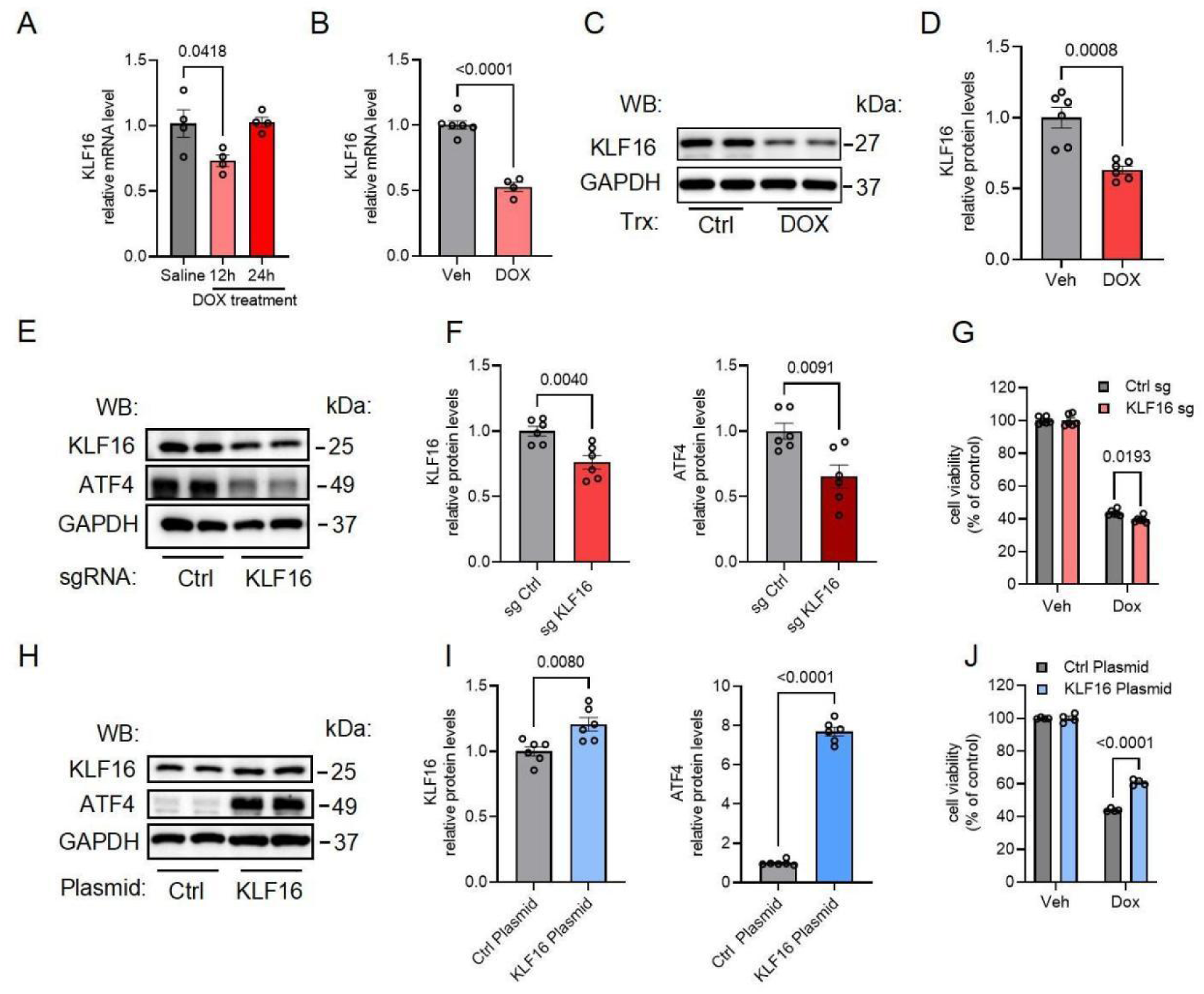
ATF4 is regulated by KLF16 instead of ISR in DIC. **(A)** RT-qPCR revealed that DOX treatment led to a decrease in KLF16 mRNA levels in the mouse heart (n = 4 mice per group). **(B)** RT-qPCR revealed that DOX treatment led to a decrease of KLF16 mRNA levels in the H9c2 cells (n = 3 per group). **(C)** Western blot revealed that DOX treatment led to a decrease in KLF16 protein levels in the H9c2 cells. **(D)** Quantification of (C) (n = 6 per group). **(E)** Relative western blots of KLF16 and ATF4 were shown in H9c2 cells transfected with control and KLF16 sgRNAs. **(F)** Quantification of (E) (n = 6 per group). **(G)** Cell viability was detected in control and KLF16 knockout cells respectively, after being treated with DOX utilizing the CCK8 assay (n = 6 per group). **(H)** Relative western blots of KLF16 and ATF4 were shown in 293T cells transfected with control and KLF16 plasmids. **(I)** Quantification of (H) (n = 6 per group). **(J)** Cell viability was detected in control and KLF16 overexpressed cells respectively, after being treated with DOX utilizing the CCK8 assay (n = 4 per group). Data are represented as mean ± SEM.

### 3.6. ATF4 protects heart from DIC via ATF4-CSE-H_2_S pathway

To further explore the mechanism of ATF4 regulated protection in DOX-induced cardiomyopathy, we revisited our published RNA sequencing dataset (GSE187005) and found that CSE was one of the top upregulated genes in ATF4 overexpressing NRVMs [14](Fig. 6A). The cysteine and methionine metabolism pathway was also enriched in the ATF4 overexpressing group(Fig. S5A-C). The significant induction of CSE by ATF4 was validated using western blotting and RT-qPCR in H9c2 cells (Fig. 6B, S5D and E). In contrast, CSE protein level significantly decreased in the ATF4^KO^ cells compared with that in the ATF4^WT^ cells (Fig. S5F and G). RT-qPCR also showed that the CSE mRNA level significantly decreased in the ATF4 knockdown cells compared with that in the control cells (Fig. S5H). We next set out to examine whether ATF4 could promote the transcriptional activity of CSE. The chromatin immunoprecipitation assay showed that ATF4 bound the promoters of CSE (Fig. 6C). We also performed a luciferase assay in Hela cells. We transfected the luciferase plasmid along with either control pGL3 or pGL3-CSE construct. Importantly, we found that ATF4 led to augmentation of the CSE promoter activity (Fig. 6D). To confirm that ATF4 deficiency-induced excessive cell death due to the ATF4-CSE-H_2_S axis, we treated ATF4^KO^ cells with H_2_S donor NaHS. The phenotype of ATF4^KO^ cells was rescued by NaHS utilizing CCK-8 assay after treated with DOX (Fig. 6E). To confirm the protective effect of ATF4 depend on the ATF4-CSE-H_2_S axis, we knockdown CSE with siRNA and then overexpressed ATF4 in H9c2 cells (Fig. S5I and J). DOX-induced cardiomyocyte damage can’t be alleviate by ATF4 in the CSE deficiency cells (Fig. 6F). We further checked H_2_S level in the ATF4 overexpressing cells utilizing SF7-AM staining. Consistently, H_2_S level significantly decreased in ATF4 overexpressing cells compared to that of the control cells after treated with DOX (Fig. 6G and H). In contrast, H_2_S level significantly decreased in ATF4 knockdown cells compared with that of the control cells after treated with DOX (Fig. 7I and J). Taken together, these data support a model that ATF4 promote the transcriptional activity of CSE and production of H_2_S, which may be one of the underlying mechanisms of ATF4-mediated cardiac protection in response to DOX-induced cardiomyopathy (Fig. 6K).

**Figure 6.**
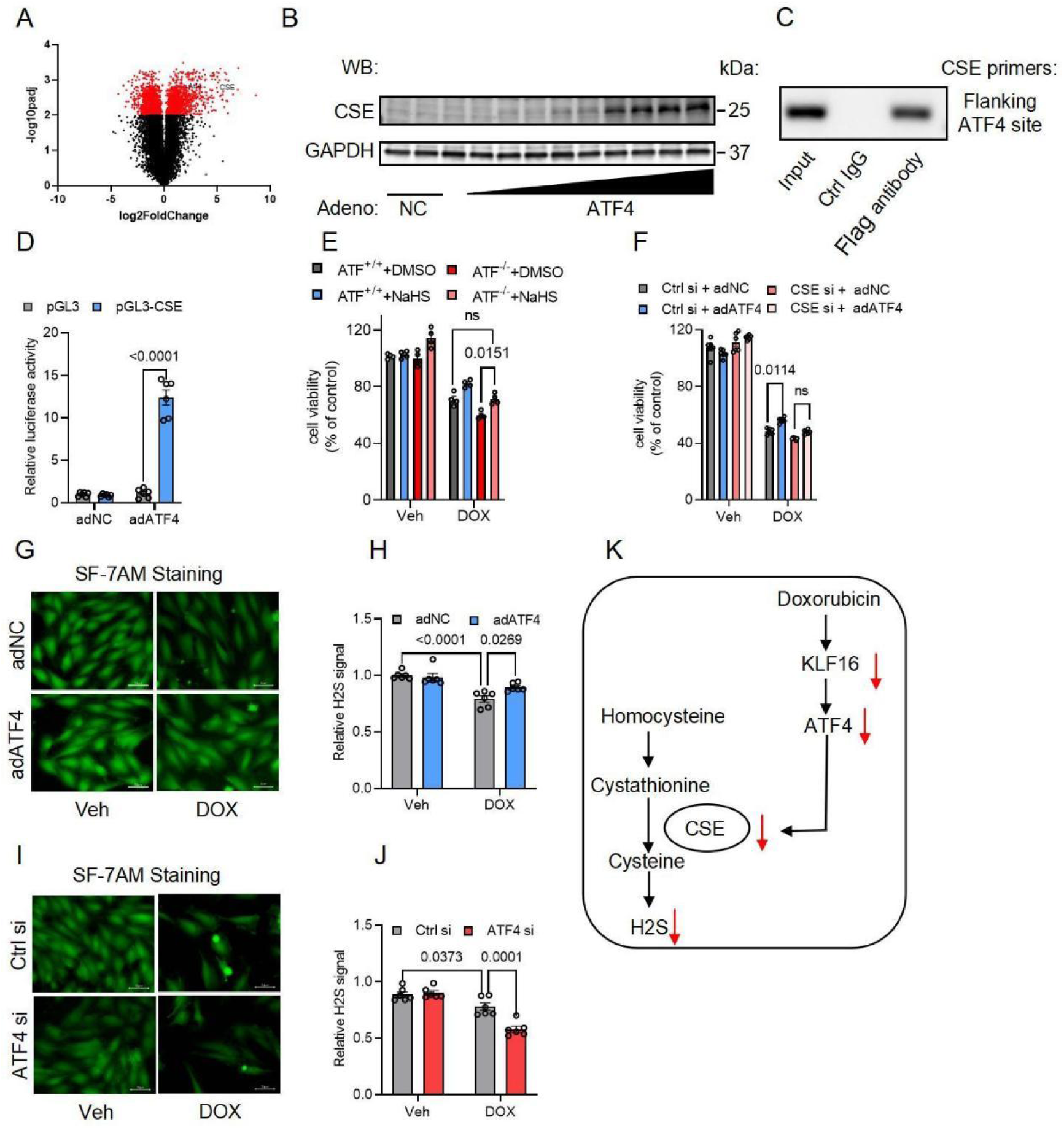
ATF4 protect heart from DIC via ATF4-CSE-H_2_S pathway. **(A)** Volcano plots detailing significant gene expression changes for adATF4 vs. adNC, with significantly changed genes in red and not significantly changed genes in black. **(B)** Relative protein levels of CSE in control and ATF4-overexpressed cells. **(C)** Chromatin immunoprecipitation (ChIP) assay showed that ATF4 bound the promoters of CSE. **(D)** Luciferase assay was performed to analyze the effect of ATF4 on CSE promoter activation (n = 6 per group). **(E)** Cell viability was detected in ATF4^+/+^ and ATF4^−/−^ cells after being treated with DOX and NaHS utilizing the CCK8 assay (n = 4 per group). **(F)** Cell viability was detected in CSE knockdown cells and control cells with infected with adNC and adATF4 after being treated with DOX utilizing the CCK8 assay (n = 4 per group). **(G)** Representative SF-7AM staining images of control and ATF4-overexpressed cells after DOX treatment were shown. **(H)** Quantification of (G). **(I)** Representative SF-7AM staining images of control and ATF4 knockdown H9c2 cells after DOX treatment were shown. **(J)** Quantification of (I). **(K)** A model of ATF4 promote the transcriptional activity of CSE and production of H_2_S. Data are represented as mean ± SEM.

**Figure 7.**
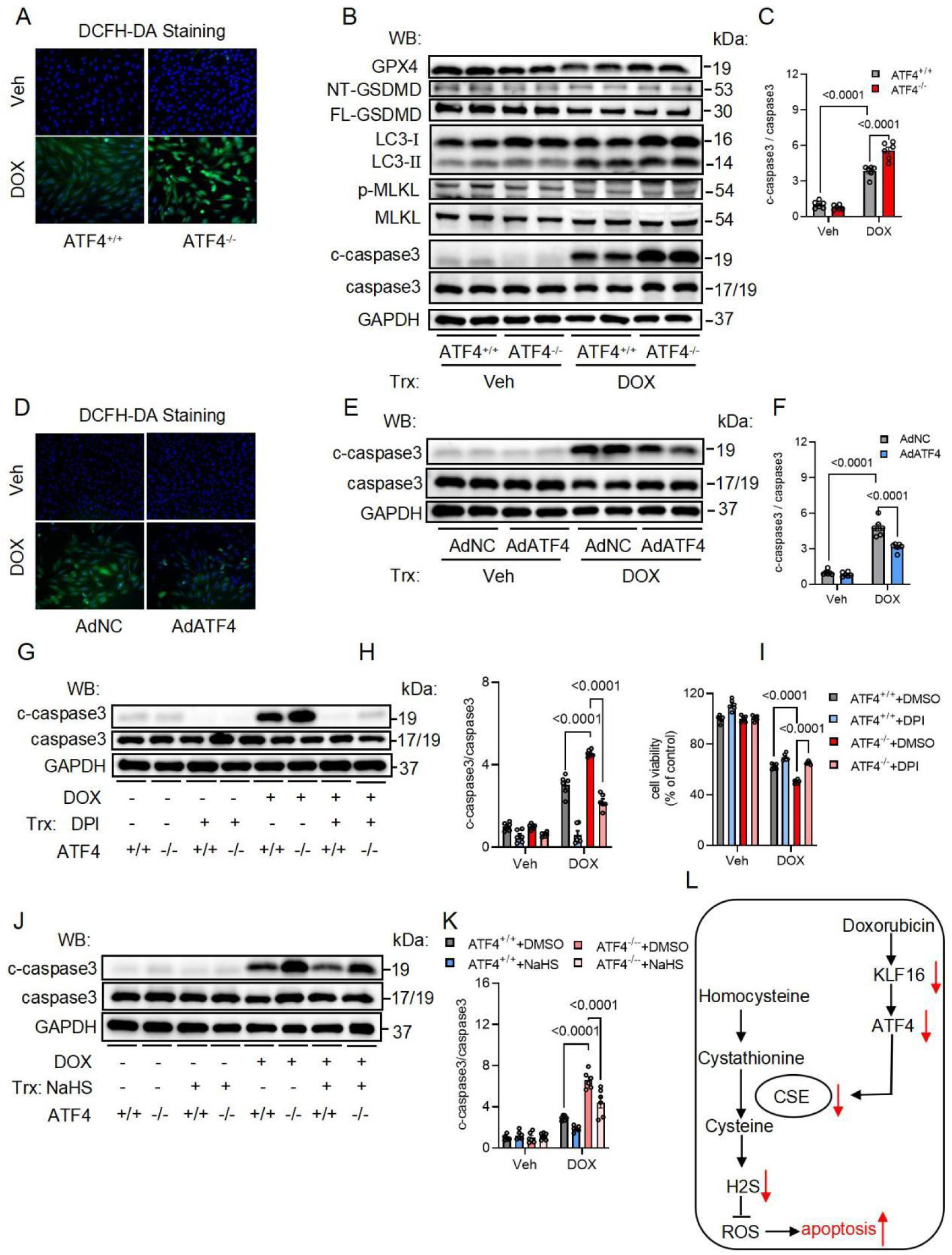
ATF4 attenuates DOX induced ROS and apoptosis. **(A)** Representative DCFH-DA staining images of ATF4^+/+^ and ATF4^−/−^ cells after DOX treatment were shown and quantified. **(B)** Relative protein levels of GPX4, GSDMD, LC3, MLKL, and Caspase3 in ATF4^+/+^ and ATF4^−/−^ cells after DOX treatment were shown. **(C)** Quantification of (B), (n = 6 per group). **(D)** Representative DCFH-DA staining images of control and ATF4-overexpressed cells after DOX treatment were shown and quantified. **(E)** Relative protein level of cleaved Caspase3 and Caspase3 in control and ATF4-overexpressed cells after DOX treatment were shown. **(F)** Quantification of (E). **(G)** Relative protein level of cleaved Caspase3 and Caspase3 in ATF4^+/+^ and ATF4^−/−^ cells after being treated with DOX and DPI. **(H)** Quantification of (G) (n = 6 per group). **(I)** Cell viability was detected in ATF4^+/+^ and ATF4^−/−^ cells after being treated with DOX and DPI utilizing the CCK8 assay (n = 6 per group). **(J)** Relative protein level of cleaved Caspase3 and Caspase3 in ATF4^+/+^ and ATF4^−/−^ cells after being treated with DOX and NaHS. **(K)** Quantification of (J) (n = 6 per group). **(L)** A model of ATF4 antagonize DOX-induced oxidative stress and apoptosis. Data are represented as mean ± SEM.

### 3.7. ATF4 attenuates DOX induced ROS and apoptosis

Since oxidative stress plays an essential role in DOX-mediated cardiomyocyte damage. We detected ROS level after DOX treatment in H9c2 cells. A marked increase of ROS level was observed in the ATF4^KO^ cells compared with that in the ATF^WT^ cells utilizing DCFH-DA staining after treated with DOX (Fig. 7A and S5K). To further explore the effect of DOX on cardiomyocyte, we checked several cell death signal pathways in DOX treated H9c2 cells. Western blotting showed that the protein levels of cleaved caspase-3 were significantly increased in the ATF4^KO^ cells compared with that in the ATF4^WT^ cells (Fig. 7B and C). To determine if ATF4 overexpression can alleviate apoptosis, we overexpressed ATF4 in H9c2 cells and treated the cells with DOX. A marked decrease of ROS level was observed in ATF4 overexpression cells compared with that in the control cells after treated with DOX (Fig. 7D and S5L). Western blotting showed that the protein level of cleaved caspase-3 significantly decreased in the ATF4 overexpression group compared with that in the control group (Fig. 7E and F). To evaluate the effect of ROS scavenger, we treat the ATF4^KO^ cells with DPI and DOX together, and we found that the phenotype of ATF^KO^ cells was rescued by DPI utilizing CCK-8 assay (Fig. 7G). Western blotting also showed that cleaved caspase-3 significantly decreased after treated with DPI (Fig. 7H and I). To evaluate the effect H_2_S donor NaHS, we treat the ATF4^KO^ cells with NaHS and DOX together. Western blotting showed that cleaved caspase-3 significantly decreased after treated with NaHS (Fig. 7J and K). Taken together, these data support a model that ATF4 promote the transcriptional activity of CSE and production of H_2_S, which can antagonize DOX-induced oxidative stress and apoptosis in cardiomyocytes(Fig. 7L).

## 4. Discussion

### ATF4 in cardiovascular disease

Emerging evidence highlights ATF4’s dual role in cardiac pathophysiology. Our previous studies showed that ATF4 antagonizes oxidative stress by regulating the production of NADPH in the cytosol and mitochondria in heart failure [14]. Similarly, ATF4 has also been shown to rescue dilated cardiomyopathy (DCM) phenotypes in induced pluripotent stem cell-derived cardiomyocytes (iPSC-CMs). This effect is attributed to the activation of the *de novo* serine biosynthesis pathway and TRIB3 kinase signaling [16]. In diabetic cardiomyopathy, ATF4 has been shown to preserve cardiomyocyte viability by upregulating FGF21-ferritin signaling under hyperglycemic stress [17]. Notably, ATF4 has been shown to play a pivotal role in cardiomyocyte regeneration, a process that involves the inhibition of mitochondrial translation via the mitochondrial stress response and the ISR-eIF2α-ATF4 axis [18]. However, excessive activation of ATF4 has been observed to induce autophagy and lethal cardiac atrophy [19]. Furthermore, the hyperactivation of ATF4 has been shown to promote the expression of TGF-β in cardiomyocytes, thereby facilitating cardiac fibrosis in the context of the DSG2^F536C^ variant of arrhythmogenic cardiomyopathy (ACM) [20]. These context-dependent roles of ATF4 position it as a critical nodal point for the fine-tuning of cardiomyocyte survival and death. Notwithstanding these advances, the involvement of ATF4 in DIC remains unexplored. Here we demonstrate a significant decrease in the expression of ATF4 in DIC. This outcome was particularly noteworthy given that earlier studies had suggested that ATF4 exhibited a modest increase in DOX-induced cardiotoxicity in diabetic rats and DOX-induced cardiac cells [21, 22]. It should be noted that the aforementioned two studies did not evaluate the ATF4 level in the DIC mouse model or the long-term treatment of DOX *in vitro*. This is the first time in which find the expression of ATF4 has been observed to decrease in DIC, a phenomenon that does not align with the established regulatory patterns of the canonical upstream regulator eIF2α. We further discovered that the decrease of ATF4 in mice exacerbated DIC, while the overexpression of ATF4 was found to be protective against this phenotype. Additional *in vitro* experiments demonstrated that the absence of ATF4 led to exacerbated DOX-mediated cardiomyocyte damage, while the overexpression of ATF4 result in a mitigation of this damage.

### KLF16 regulate ATF4 in DIC

DIC can manifest as hypotension, tachycardia, arrhythmias, and ultimately congestive heart failure [23]. The precise mechanism of DIC remains to be elucidated. However, it is well established that the production of oxygen free radicals plays an instrumental role in its progression. ATF4 is a pivotal transcription factor that has been demonstrated to promote the expression of a variety of cytoprotective genes. These genes associated with amino acid metabolism, oxidative stress damage, and ER-phagy in response to cellular stress [24, 25]. This study indicates that a reduction of ATF4 exacerbate DIC which is validated by cardiac function and survival outcomes *in vivo*. Consistently, the *in vitro* data demonstrated that ATF4 deficiency results in increased DOX-mediated cardiomyocyte damage. This finding is corroborated by cell viability testing and Sytox Green staining. ATF4 is the know downstream component of eIF2α, which is the core of the ISR [26]. Contrary to our initial hypothesis, we observed a substantial increase in the phosphorylation level of eIF2α, rather than the anticipated decrease. We then sought to identify other upstream regulators of ATF4 in the context of DIC. We demonstrated a decline in KLF16 mRNA and protein levels in DIC. Moreover, ATF4 protein levels significantly decreased following KIF16 knockdown. Consistently, KIF16 knockdown exacerbated DOX-mediated cardiomyocyte damage. Conversely, ATF4 protein levels exhibited a substantial increase in response to KIF16 overexpression. Consistently, KIF16 overexpression alleviated DOX-mediated cardiomyocyte damage.

The KLF family contains 17 members, the majority of which play a role in embryogenesis, development, and homeostasis [27]. A substantial body of research has indicated that KLFs serve as either tumor suppressors or oncogenes, depending on the cellular contexts [28, 29]. KLF16 has been demonstrated to play a pivotal role in the metabolism and regulation of the endocrine system [30, 31]. A hepatic KLF16 knockout mouse model revealed that KLF16 is associated with hepatic lipid homeostasis and redox homeostasis by regulating the transcription of PPARα [32]. In this study, we demonstrate that KLF16 modulates the transcription of ATF4, thereby counteracting oxidative stress in DOX-mediated cardiomyocyte damage.

### ATF4 is the upstream regulator of H_2_S production

The endogenously produced H_2_S is derived from L-cysteine through the actions of three enzymes: cystathionine β-synthase (CBS), cystathionine γ-lyase (CSE), and 3-mercaptopyruvate sulfurtransferase (3-MST) [33]. H_2_S has been classified as a toxic gas due to its ability to impede the activity of cytochrome c oxidase and disrupt mitochondrial respiration. However, it is important to note that varying exposure concentrations of H_2_S may yield different biological effects. A substantial body of evidence has emerged indicating that a minimal level of H_2_S is produced endogenously in mammalian cells and exerts various biological functions. It has been demonstrated to combat oxidative species, react with the metal centers of iron-heme proteins, and modify protein cysteine residues known as persulfidation [34, 35]. The interaction of H_2_S with various signaling molecules has been postulated to facilitate signaling transduction and play a pivotal role in cardiovascular diseases, including atherosclerosis, coronary heart disease, heart failure, hypertension, pulmonary arterial hypertension, and myocardial ischemia/reperfusion injury [36–38]. In the context of DIC, H_2_S has also been demonstrated to exert a protective effect [39–41]. However, the upstream regulator of H_2_S in DIC remains to be elucidated. Our findings identified ATF4, a well-established mediator of the integrated stress response, as a critical regulator of CSE and H_2_S production in DIC. A subsequent investigation of the RNA-seq data set revealed that CSE was ranked within the top 5 genes that demonstrated a significant increase in expression. This finding was corroborated *in vitro* under ATF4 gain-and-loss of function conditions. Furthermore, an increase in H_2_S levels was observed in the condition of ATF4 overexpression in H9c2 cells. Our findings on the cardioprotection by ATF4 support a prevailing notion that ATF4 exerts a beneficial effect by antagonizing oxidative stress under DIC conditions.

### Conclusions and Perspectives

The predominant pathogenesis of DIC is ROS production. Our previous study indicates that ATF4 can antagonize oxidative stress in heart failure under pressure overload. The present study demonstrated that ATF4 is potently suppressed in DIC, which exacerbates the cardiac damage due to decreased H_2_S levels and increased ROS production. The restoration of ATF4 mitigates cardiac damage in DIC by augmenting H_2_S levels and diminishing ROS production. Importantly, our findings revealed that the expression level of ATF4 was subject to regulation by KLF16, rather than the conventional upstream regulator eIF2α, in the context of DOX-induced cardiomyopathy. In summary, our findings reveal a critical role of the KLF-ATF4-H_2_S in DOX-induced cardiomyopathy, which may advance our understanding of DIC pathophysiology and provide a promising target to curb DIC.

## Author Contributions

**Shuting Xu:** Writing – review & editing, Writing – original draft, Visualization, Validation, Project administration, Methodology, Investigation, Formal analysis, Data curation. **Yan Shi:** Methodology, Investigation, Formal analysis, Data curation. **Xiaoshuai Zhao:** Methodology, Investigation, Formal analysis. **Xuhui Chen:** Methodology, Investigation, Formal analysis. **Ying Liu:** Resources. **Fan Zhang:** Methodology. **Fangchi Yu:** Investigation. **Linhao Ruan:** Resources. **Chaolong Wang:** Methodology. **Xuejun Jiang:** Writing – review & editing, Supervision. **Xiaoding Wang:** Writing – review & editing, Writing–original draft, Supervision, Resources, Project administration, Conceptualization. **Guangyu Zhang:** Writing – review & editing, Writing – original draft, Visualization, Validation, Project administration, Methodology, Investigation, Funding acquisition, Formal analysis, Data curation, Conceptualization.

## Data availability statement

The data related to this article are incorporated into the article and its supplementary material. Raw bulk RNA sequencing data have been deposited at the National Center for Biotechnology Information Gene Expression Omnibus (GSE187005).

## Ethics approval statement

All animal experimental procedures were approved by the Biomedical Animal Ethics Committee of Health Science Center, Tongji Hospital carried out following their legal requirements (No. T1-202505038).

## Acknowledgments

We thank Christopher M. Adams (Division of Endocrinology, Metabolism and Nutrition, Department of Medicine, Mayo Clinic, Rochester, Minnesota) for kindly providing ATF4^F/F^ mice.

## Footnote

## Nonstandard Abbreviations and Acronyms

DOX: Doxorubicin
ROS: reactive oxygen species
ATF4: Activating transcription factor 4
DIC: DOX-induced cardiomyopathy
CSE: cystathionine γ-lyase
H_2_S: hydrogen sulfide
ISR: integrated stress response
elF2a: α-subunit of eukaryotic initiation factor 2
ER: endoplasmic reticulum
TFs: transcription factors
KLFs: Krüppel-like factors
DCM: dilated cardiomyopathy
iPSC-CMs: induced pluripotent stem cell-derived cardiomyocytes
ACM: arrhythmogenic cardiomyopathy
CBS: cystathionine β-synthase
3-MST: 3-mercaptopyruvate sulfurtransferase

